# The balance between antiviral and antibacterial responses during *M. tuberculosis* infection is regulated by the ubiquitin ligase CBL

**DOI:** 10.1101/2024.05.15.594178

**Authors:** Tina Truong, Kelsey Martin, Michelle Salemi, Abigail Ray, Brett S. Phinney, Bennett H. Penn

**Affiliations:** Department of Internal Medicine, University of California, Davis, Davis, California, United States of America; Graduate Group in Immunology, University of California, Davis, Davis, California, United States of America; Proteomics Core Facility, University of California, Davis, Davis, California, United States of America; Microbiology Graduate Group, University of California, Davis, Davis, California, United States of America; Department of Medical Microbiology and Immunology, University of California, Davis, Davis, California, United States of America

## Abstract

As a first line of host defense, macrophages must be able to effectively sense and respond to diverse types of pathogens, and while a particular type of immune response may be beneficial in some circumstances, it can be detrimental in others. Upon infecting a macrophage, *M. tuberculosis* (*Mtb*) induces proinflammatory cytokines that activate antibacterial responses. Surprisingly, *Mtb* also triggers antiviral responses that actually hinder the ability of macrophages to control *Mtb* infection. The ubiquitin ligase CBL suppresses these antiviral responses and shifts macrophages toward a more antibacterial state during *Mtb* infection, however, the mechanisms by which CBL regulates immune signaling are unknown. We found that CBL controls responses to multiple stimuli and broadly suppresses the expression of antiviral effector genes. We then used mass-spectrometry to investigate potential CBL substrates and identified over 46,000 ubiquitylated peptides in *Mtb*-infected macrophages, as well as roughly 400 peptides with CBL-dependent ubiquitylation. We then performed genetic interaction analysis of CBL and its putative substrates, and identified the Fas associated factor 2 (FAF2) adapter protein as a key signaling molecule protein downstream of CBL. Together, these analyses identify thousands of new ubiquitin-mediated signaling events during the innate immune response and reveal an important new regulatory hub in this response.

## Introduction

Being among the first cells to respond to infection, tissue macrophages must be able to sense and effectively respond to diverse types of pathogens, ranging from multicellular eukaryotic parasites to bacteria and viruses. Longstanding clinical observations have suggested potential antagonism between immune responses targeting different classes of microbes. Patients with preceding influenza virus infection carry roughly a 100-fold higher near-term risk of developing bacterial pneumonia from *Streptococcus pneumoniae* (1). Similarly, while CD4 T helper 2 (Th2) cells are critical for responses to multicellular parasites, patients with Th2-dominant immune responses in *Mycobacterium leprae* lesions harbor higher bacterial loads and experience more severe disease (2).

Even in the simple scenario of a single immune cell encountering a lone pathogen, the mechanisms by which the type of pathogen is discerned, and an appropriate response mounted remains poorly understood. Dozens of individual sensors for diverse bacterial-, fungal-, and virus-associated PAMPS have been described (3,4). However, many pathogens, including *M. tuberculosis* (*Mtb*), activate multiple sensors simultaneously. How macrophages integrate multiple, sometimes discordant, pathogen-associated signals into a coherent, pathogen-appropriate, response remains unclear.

*Mtb* persists as a threat to human health. It currently infects one-third of the world’s population, causing a chronic, often lifelong infection, and tuberculosis (TB) kills an estimated 1-2 million people annually (5). Macrophages infected with *Mtb* are immediately exposed to an array of cell wall-derived pathogen-associated molecular patterns (PAMPs) such as lipoproteins, peptidoglycan and trehalose 6,6’-dimycolate, which activate cognate receptors such as TLR2, NOD2 and CLEC4E (Mincle) (6–14). The outcome of many of these signaling pathways is the activation of nuclear factor kappa B (NFκB) family transcription factors and the upregulation of proinflammatory cytokines such as TNF and IL1B which activate potent antibacterial responses and are critical for controlling *Mtb* replication (15–22).

Surprisingly, *Mtb* also activates numerous antiviral effectors in infected macrophages. *Mtb* rapidly permeabilizes its phagosome and releases bacterial nucleic acids into the host cytosol through unclear mechanisms (24–26). Bacterial DNA is recognized by cyclic GMP-AMP synthase (cGAS) which then triggers a cascading activation of STING1, TBK1 and the transcription factors IRF3 and IRF7. These induce hundreds of genes, including the type I interferons like interferon beta (IFN-β) that propagate the signal to nearby cells (24–31). Bacterial RNA is also introduced into the host cell, initiating signaling through the retinoic acid-Inducible gene I (RIG-I) sensor which triggers MAVS-dependent activation of IRF3/IRF7 with similar effects (25). While these initial observations came from mouse models, gene expression analysis of TB patients has detected similar activation of type I IFN in the peripheral blood of patients with active TB (32–34).

Several lines of evidence demonstrate that these antiviral responses interfere with the antibacterial capacity of macrophages, and suggest that macrophages can be polarized into antiviral or antibacterial states by their encounter with a pathogen. C3H-derived strains of mice harbor a polymorphism causing loss of the SP140 transcriptional repressor, resulting in hyperactivation IFN-β. These strains are highly susceptible to *Mtb*, succumbing rapidly with dramatically increased bacterial burdens (35–39) – a phenotype that is rescued by the administration of an IFN-β blocking antibody or by loss of the type I interferon receptor (IFNAR) (39,40). C57BL/6 mice infected with *Mtb* produce lower levels of IFN-β following infection, but even in this context, loss of MAVS or IFNAR results in decreased bacterial burden and increased survival of infected animals (25,26) Thus, activation of antiviral responses within the host seems to actively interfere with an effective antibacterial response to *Mtb*, and suggests that *Mtb* might introduce nucleic acids into the host cell as a virulence strategy,

Ubiquitin signaling plays an important role in regulating both antibacterial and antiviral responses in *Mtb*-infected macrophages. Ubiquitin ligases such as PARKIN and SMURF are critical in targeting *Mtb* for autophagy (41–44). Separately, the ubiquitin ligases TRIM14 and CBL have been shown to regulate the expression of type I interferon following *Mtb* infection (45,46). CBL is targeted by the secreted *Mtb* virulence factor LpqN, which is required for *Mtb* replication in mice (45). Proteomic analyses identified a physical interaction between LpqN and CBL, and the growth of the attenuated *lpqN Mtb* mutant was rescued by disruption of host *Cbl*. While *Cbl*-deficient macrophages are more permissive for *Mtb* replication, the opposite is seen with viral challenge, as CBL-deficient macrophages more potently restrict viral replication (45). Together, these findings suggest that CBL is an important regulator of the balance between antiviral versus antibacterial responses in macrophages. However, how CBL regulates this process is not known.

Here, we investigated the mechanisms by which CBL regulates immune signaling. We found that it controls macrophage responses to multiple stimuli and represses the expression of numerous antiviral effectors. We then used proteomics to identify potential CBL substrates in macrophages as they respond to *Mtb* infection and identified the endoplasmic reticulum-localized signaling adapter Fas-associated factor 2 (FAF2) as a protein that undergoes CBL-dependent ubiquitylation. Notably, while CBL deficient macrophages are more permissive for *Mtb* growth, simultaneous loss of FAF2 in double-knockout macrophages reverses this phenotype, suggesting that the FAF2 pathway might be hijacked by *Mtb,* and that CBL acts to constrain FAF2 activity in *Mtb*-infected macrophages.

## Results

### CBL broadly suppresses antiviral effector expression

Prior work has shown that CBL suppressed host antiviral responses in mouse primary bone-marrow-derived macrophages (BMDMs) following *Mtb* infection (45). We began by assessing whether this role of CBL was evolutionarily conserved, with CBL also regulating these processes in human cells. We used shRNA to deplete CBL mRNA in the human THP-1 cell line that differentiates into macrophage-like cells after stimulation with phorbol-12-myristate-13-acetate (PMA) and 1,25-dihydroxy-vitamin D, and found an ∼85% reduction in CBL mRNA levels (Fig 1A). We first assessed whether loss of CBL in human macrophages would similarly rescue the growth of the CBL-sensitive *lpqN* mutant *Mtb* strain which lacks the effector that normally antagonizes CBL function, and which is attenuated in both mouse BMDMs and in mice (45). Following differentiation, we infected THP-1 control cells expressing a non-targeting shRNA, and CBL-depleted cells expressing a CBL-specific shRNA. To monitor bacterial replication, we used the *lpqN Mtb* mutant carrying a *LuxBCADE* bioluminescent reporter operon (45), and monitored bacterial growth over time by quantifying luminescence. Consistent with prior observations in mouse BMDMs, we found that loss of CBL also rescued the growth of the *lpqN* mutant in human macrophages (Fig 1B). The increased bacterial replication was confirmed by direct colony forming unit (CFU) numeration (Fig S1A) and was associated with a more rapid loss of macrophage viability in CBL-deficient macrophages over the course of infection (Fig 1C). The effect of CBL on *Mtb* replication was independently verified using a second independent shRNA in THP-1 cells (Fig S1B). To confirm these findings, we also evaluated the role of CBL in a second human macrophage line, HL-60. Using shRNA to deplete CBL in these cells (Fig 1D) we found that loss of CBL similarly rescued the growth of the *lpqN* mutant in HL-60 cells as well (Fig 1E).

**Fig 1.**
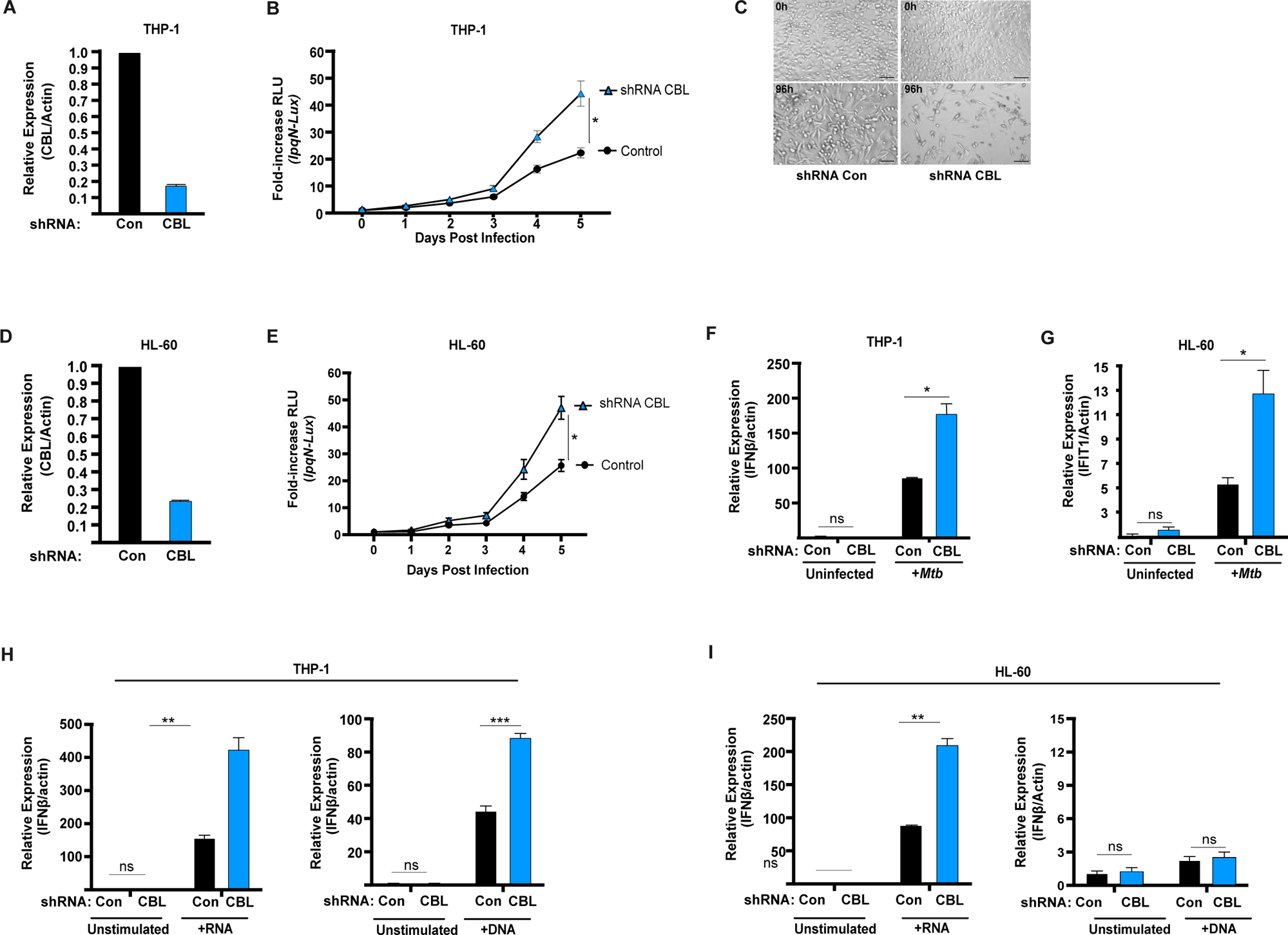
CBL suppresses antiviral responses in human cells. (A) CBL expression analyzed by RT-qPCR analysis of THP-1 cells expressing CBL-specific shRNA or a control non-targeting shRNA. (B) Luminescent growth assay of CBL-sensitive *lpqN Mtb* mutant carrying the *LuxBCADE* operon. (C) Phase-contrast microscopy images of the same monolayer of THP-1 macrophages prior to infection and 96 h post-infection by *lpqN Mtb* (MOI=0.8, 10x magnification; scale bar 100μm). (D) RT-qPCR analysis of CBL expression in HL-60 cells. (E) Luminescent growth assay of *lpqN Mtb* in HL-60 cells. (F),(G) RT-qPCR analysis of *lpqN Mtb*-infected macrophages 6 h after infection. Analysis of (H) THP-1 or (I) HL60 cells transfected with either RNA or DNA as indicated. Error bars denote SEM of technical replicates; statistical significance was evaluated by two-tailed t-test and indicated on plot: *p≤0.05, **p≤0.01, ***p≤0.001, ns p>0.05. Representative (median) data of three or more independent experiments are shown.

We next determined whether expression of antiviral effectors was also regulated by CBL in human cells. Consistent with prior observations in mouse macrophages, we found that depletion of CBL resulted in an increased expression of antiviral effectors in both THP-1 and HL-60 cells following *Mtb* infection (Fig 1F,G). We next used defined PAMPs to selectively stimulate different pathways leading to IFN-β activation to assess if CBL was regulating a single sensor pathway or multiple pathways. Work from several groups has shown that *Mtb* activates IFN-β both through the cGAS DNA sensing pathway as well as the RIG-I RNA sensing pathway (24–25,31–32). To assess which pathways are regulated by CBL, we individually transfected THP-1 and HL-60 macrophages with either dsDNA, a known agonist for cGAS, or with 5′-phosphorylated RNA, a known agonist for RIG-I, in order to evaluate the role of CBL. We found that in response to either DNA or 5’ phosphorylated RNA, loss of CBL resulted in an increased expression of IFN-β in THP-1 cells (Fig 1H). In HL-60 cells we observed a similar role for CBL in regulating RNA-stimulated IFN-β responses. However, following DNA stimulation HL-60 cells displayed only weak induction of IFN-β, and did so independently of CBL (Fig 1I). Taken together these data support a model where the role of CBL in innate immunity is evolutionarily conserved, acting to suppress expression of antiviral effectors such as IFN-β in multiple different contexts.

We next sought to determine whether CBL selectively regulated a restricted group of genes, such as IFIT1 and IFN-β, or whether it orchestrated broad transcriptional changes in macrophages as they responded to *Mtb*. To assess this, we infected THP-1 cells with *Mtb*, using the *lpqN* mutant strain so that CBL activity would be unconstrained by the bacterial effector. We infected both control and CBL-deficient THP-1 cells for 6 h, a time at which many immune-related genes are strongly induced, and prepared RNA for expression profiling by RNA-Seq. As expected, *Mtb* infection of control cells triggered widespread gene expression changes with upregulation of 735 genes and downregulation of 423 genes (FDR≤0.05, Log2 fold-change>0.5) and strong induction of both proinflammatory and antiviral effectors. (Fig S2A, Table S1). CBL-deficient cells regulated most of these genes in a similar manner following *Mtb* infection. However, a set of genes showed distinct CBL-dependent transcriptional changes, with 204 genes hyperactivated and 62 genes depressed in CBL-deficient cells following *Mtb* infection (Fig 2A and Table S1). Interestingly, while CBL regulates the production of IFN-β in these cells (Fig 1G,I) it does not regulate their response to IFN-β stimulation (Fig S2B), suggesting that the broad CBL-dependent gene expression changes are due to cell-intrinsic mechanisms rather than the indirect effects of type I interferon.

**Fig 2.**
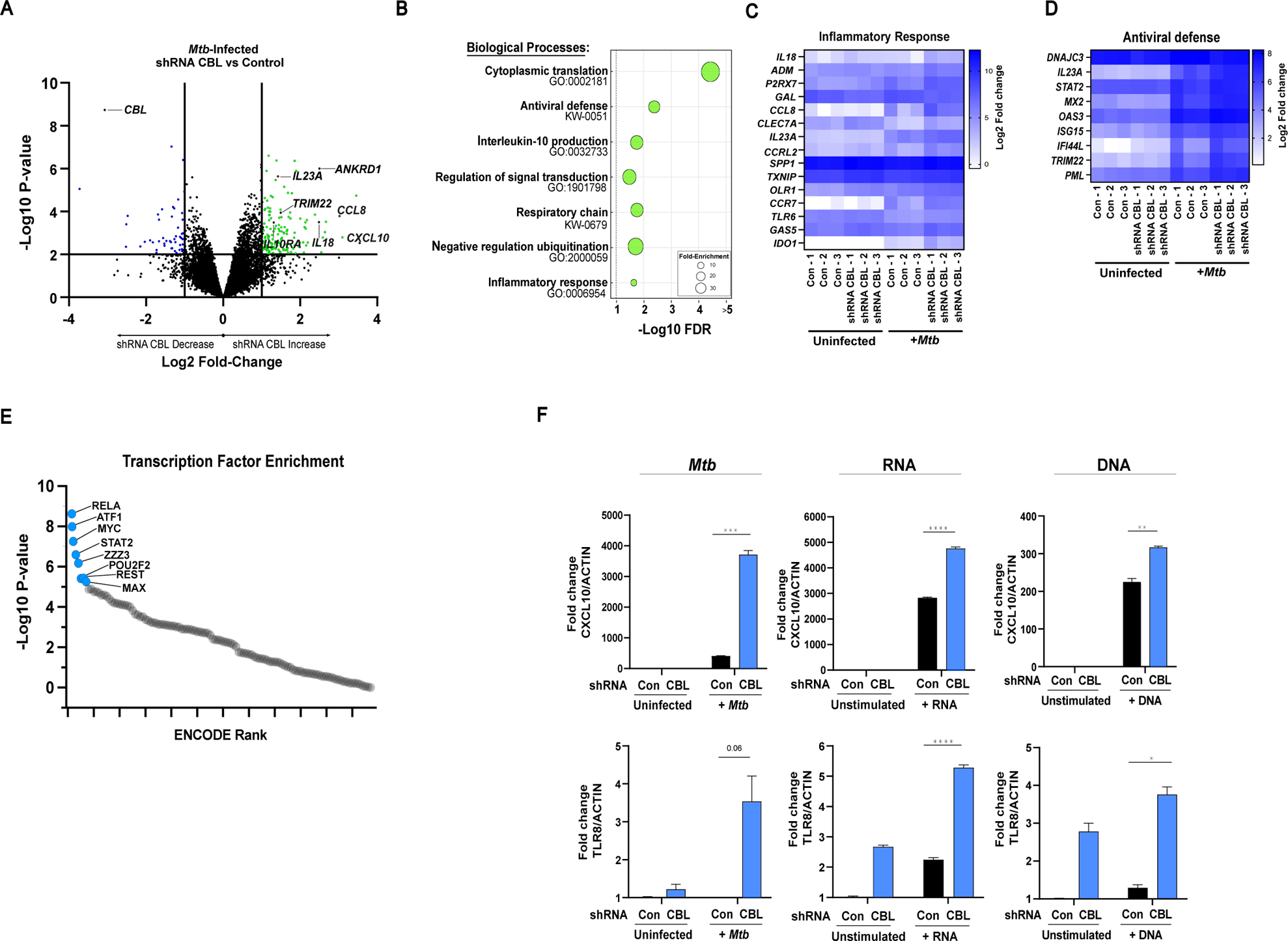
CBL regulates a complex transcriptional program. (A) RNA-Seq gene expression analysis of THP-1 macrophages expressing CBL-specific shRNA vs control non-targeting shRNA 6 h after infection with *lqpN*-mutant *Mtb*. (B) Biological processes transcriptionally regulated by CBL. (C), (D) Differential expression of immunity-related genes in CBL-deficient cells. (E) ENCODE Transcription factor binding site enrichment of CBL-regulated genes (F) RT-qPCR analysis of CBL-regulated genes after indicated stimuli. Differentially-expressed genes for analyses in (B-E) defined by RNA-Seq FDR<0.05 and Log2 fold-change>0.5. RNA-Seq analysis was performed on 3 independent experiments. For RT-qPCR, error bars denote SEM of technical replicates; statistical significance assessed with two-tailed t-test and indicated on plot. Representative (median) data of three independent experiments are shown.

We further analyzed the set of genes regulated by CBL to evaluate which cellular processes were being affected, and what transcription factors might be mediating the regulatory effects of CBL. We performed systematic pathway enrichment analysis using the Gene Ontology and KEGG Databases and identified several clusters of functionally-related genes regulated by CBL. Supporting the hypothesis that CBL broadly suppresses antiviral responses, we found a significant over-representation of antiviral defense genes in the set of CBL-regulated genes (10-fold over-representation, FDR <0.02). We also found significant associations with several other pathways including IL-10 production and global protein translation (Fig 2B-D). In addition, we analyzed the set of genes transcriptionally regulated by CBL to determine whether any associations existed between these genes and characterized transcriptional regulators. We assessed this by systematically querying the ENCODE Chip-Seq database (47) with the set of genes regulated by CBL. In uninfected cells, there were no transcriptional regulators enriched at these genes. In contrast, in *Mtb*-infected cells, analysis of the 266 CBL-responsive genes using ENCODE showed an association with several transcriptional regulators (Fig 2E). The most significant association was seen with for the NFkB family member RELA, with an odds-ratio (OR) of 2.6 (p≤1E-9) found at a set of immune-related genes, with additional associations noted for STAT2 (OR=5.3, p≤2E-7), ATF1 (OR=2.5, p≤1.1E-8), and MYC (OR=2.1, p≤5E-8), suggesting that these transcription factors might be mediating the transcriptional effects of CBL.

We also assessed whether genes we identified as regulated by CBL during *Mtb* infection were also regulated by CBL in response to other stimuli. We selected TLR8 and CXCL10, two genes identified by our RNA-Seq analysis of *Mtb*-infected macrophages whose expression was suppressed by CBL, and examined their regulation in cells exposed other PAMPs - either cytoplasmic dsDNA to activate cGAS or 5’-phosphorylated RNA to activate RIG-I. In both cases we found hyperactivation of these genes in CBL-deficient cells, although the patterns differed somewhat. For TLR8 we detected basal de-repression in unstimulated cells and hyperactivation following Mtb infection and either DNA or RNA stimulation. For CXCL10, there was no basal de-repression, and while CXCL10 was strongly regulated by CBL during *Mtb* infection, it showed only modest increases in expression in CBL-deficient cells stimulated with DNA or RNA (Fig 2F). Taken together, these results demonstrate that CBL regulates a broad subset of antiviral response genes in macrophages, along with regulation of several other processes, and that it does so in response to multiple stimuli.

### CBL enzymatic activity is required to regulate macrophage responses to Mtb

We next sought to determine the mechanisms by which CBL regulates immune responses. CBL is a RING-domain E3 ligase that was initially characterized as a proto-oncogene, ubiquitylating activated receptor tyrosine kinases to target them for degradation (48–50). Subsequent studies using an enzymatically inactive CBL point mutant showed that CBL also has important non-enzymatic functions, and can act as a signaling scaffold that potentiates phosphatidylinositol 3-kinase and SRC-family kinase activity (51). This non-enzymatic function plays an important role in vivo, as several of the developmental defects *Cbl^−/−^* mice could be rescued by expression of enzymatically-inactive CBL (52).

To test whether CBL was acting enzymatically to ubiquitylate key substrates during innate immune responses, or whether it was acting as a signaling adapter protein, we performed genetic complementation analysis in CBL-deficient macrophages, re-introducing either wild type CBL or catalytically inactive mutants (Fig 3A). To study human macrophages, we used shRNA-expressing CBL-deficient THP-1 cells. To study mouse macrophages we used Cas9-expressing <u>c</u>onditionally-immortalized <u>m</u>acrophages (CIMs) which are mouse myeloid precursor cells that carry a HOXB8-ER estrogen-regulatable transcription factor. This maintains the cells as immortalized precursors in the presence of estrogen, but upon estrogen withdrawal allows differentiation into macrophages closely recapitulating the physiology of primary mouse BMDMs (53). These cells display high-efficiency genome editing, and transduction of CIM cells with a CBL-specific sgRNA led to a >95% reduction in CBL protein levels in the population (Fig 3B).

**Fig 3.**
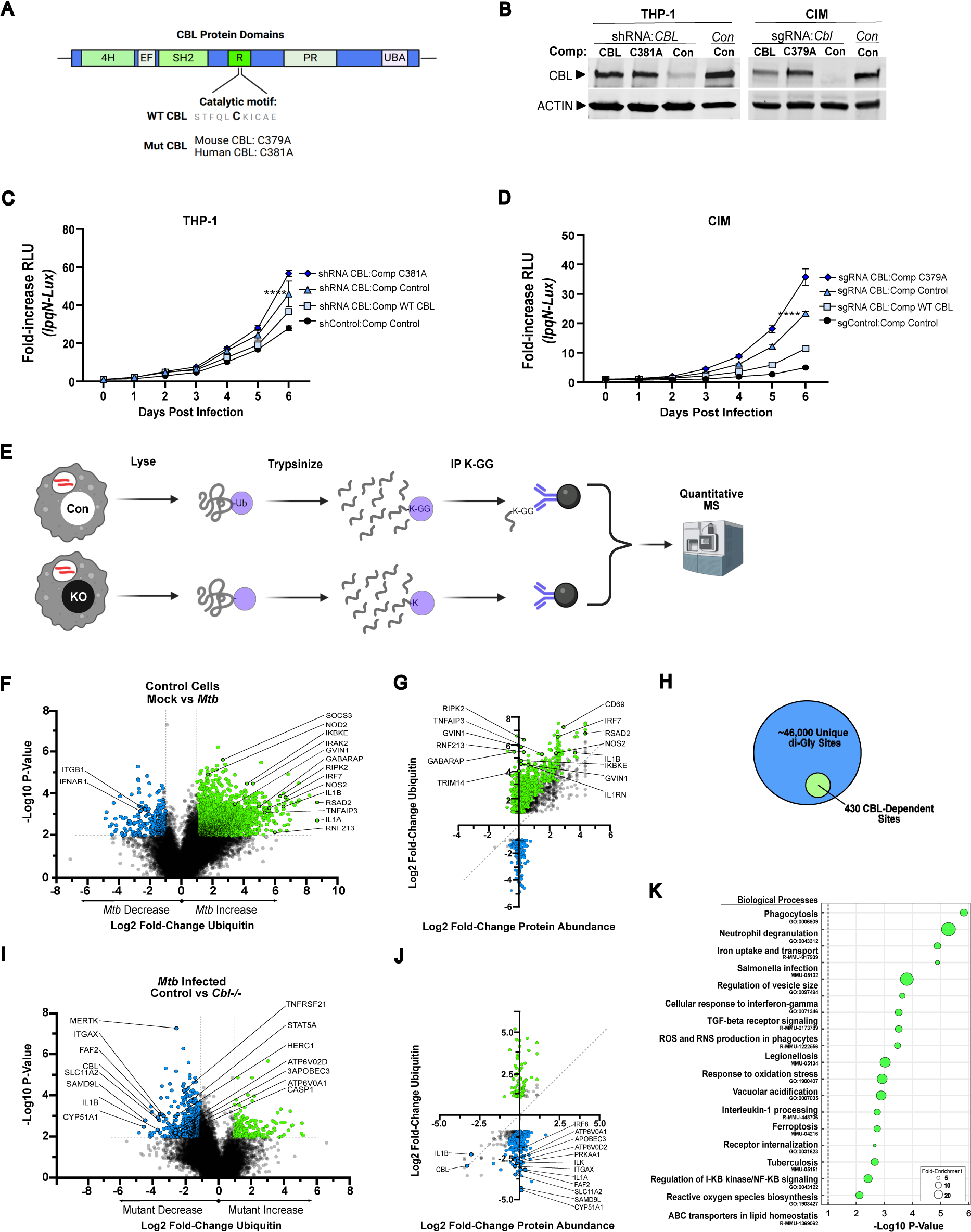
CBL-dependent ubiquitylation is required to restrict *Mtb* growth in mouse and human macrophages. (A) CBL domain structure and catalytic site mutants. (B) Immunoblot of wild-type and mutant CBL isoforms expressed from lentiviral vectors in CBL-deficient human (THP-1) and mouse (CIM) macrophages. (C), (D) Luminescent growth assay of *lpqN Mtb* in CBL-deficient macrophages in which CBL expression was restored with either wild-type or mutant CBL. (E) Experimental design for ubiquitylation proteomics using di-Gly MS. (F) di-Gly-MS analysis of ubiquitylation in control CIM cells expressing a non-targeting sgRNA 6 h post-infection with the *lpqN Mtb* mutant. Green (increased) and blue (decreased) indicate peptides with p-value ≤0.01 and Log2 fold-change >2. (G) Comparison of ubiquitylation versus total-protein changes in control cells following infection. Green (increased) and blue (decreased) denote data points where the change in di-Gly peptide abundance was >3 standard-deviations greater than the change in protein abundance. (H) Subset of CBL-dependent changes (I) di-Gly MS analysis of ubiquitylation 6 h post-infection in control vs. *Cbl*^−/−^ cells infected with *lpqN Mtb*. (J) Ubiquitylation versus total-protein changes in control vs. *Cbl*^−/−^ cells infected with *lpqN Mtb.* (K) Biological process of proteins with CBL-dependent ubiquitylation. MOI=0.8 for growth assays, and MOI=10 for di-Gly-MS. Di-gly MS was used to analyze 3 independent experiments. For growth assays, representative (median) data of three or more independent experiments are shown; p-value determined by two-tailed t-test and error bars represent SEM of technical replicates.

For both mouse and human cells, we used lentiviral vectors to deliver either catalytically inactive CBL (human C381A, mouse C379A) or WT CBL. In both cases, CBL expression was restored to ∼50% of endogenous levels, without evidence of destabilization of the mutant protein (Fig 3B). As seen previously (45), CBL-deficient macrophages were more permissive for *Mtb* replication than control cells, and as expected, in both mouse and human macrophages, re-introduction of WT CBL significantly restricted the growth of *Mtb* (Fig 3C,D). In contrast, re-introduction of catalytically-inactive CBL failed to complement the defective host response, demonstrating that enzymatic activity is needed for CBL function in this context. Interestingly cells expressing inactive CBL actually had higher bacterial loads than CBL-deficient cells, suggesting that catalytically-inactive CBL might act as a dominant-negative isoform, perhaps by interfering with the activity of other signaling molecules such as the related CBL-family ligase CBLB.

### Identifying potential CBL substrates

Given that the enzymatic activity of CBL was necessary to regulate immune signaling, we next sought to identify CBL substrates in this context. Several growth factor receptors including EGFR, MET, and CSF1R have been characterized as CBL substrates in other settings (48–50). However, the CBL substrates during innate immune responses are not known. We used *Cbl^−/−^* CIM cells to study ubiquitylation because of the near-complete loss of CBL protein in the polyclonal population of cells expressing a CBL-specific sgRNA, and their larger *Mtb* growth phenotype relative to human cells partially depleted of CBL via shRNA (Fig 3B-D). Using the *lpqN* mutant strain that is unable to antagonize CBL activity, we infected cells for 6 h, and added the proteasome inhibitor bortezomib for the final 2 h to stabilize ubiquitylated proteins and facilitate their detection.

We then used di-Gly enrichment and quantitative mass-spectrometry (di-Gly MS) to globally monitor changes in ubiquitylation in *Mtb*-infected cells (Fig 3E). Lysates were prepared from uninfected and infected cells, and from control and *Cbl^−/−^* cells, and digested with trypsin. We then enriched ubiquitylated peptides by immunoprecipitation with an antibody that recognizes the di-Gly remnant of ubiquitin that remains conjugated to the ubiquitin-modified Lys residue following trypsinization, and analyzed the samples by liquid chromatography tandem MS (LC-MS/MS), using label-free quantification of MS2 spectra (54–59). Of note, the ubiquitin-like proteins ISG15 and NEDD8 leave an indistinguishable di-Gly Lys remnant. However, these modifications account for only ∼5% of identified di-Gly modified residues, and the vast majority of di-Gly sites are ubiquitin-modified (60–62). In addition, since CBL is a ubiquitin-specific ligase, any CBL-dependent di-Gly modifications should represent CBL-dependent ubiquitylation. Thus, we will hereafter refer to di-Gly-modified sites as ubiquitylation, recognizing that a small percentage of sites modified ligases other than CBL are modified by ISG15 or NEDD8.

Overall, we identified ∼76,000 unique peptides in macrophages, encompassing ∼46,000 unique di-Gly-modified sites. This dataset creates the most detailed atlas of ubiquitin modification in macrophages to date, with roughly 10-fold more sites identified than currently available datasets of this specialized cell type. As expected, in control cells, we saw numerous ubiquitin changes in response to *Mtb* with a total of 1,793 ubiquitylation sites increasing (Log2 fold-change >1, FDR <0.05) and 197 sites decreasing (Fig 3F, Table S2). Since the quantity of a ubiquitylated peptide in a sample can change because of either a change in the stoichiometry of ubiquitylation at a given site, or because of a change in abundance of the protein itself, we sought to deconvolute these processes. We examined changes in protein abundance in the same samples by quantitative LC-MS/MS (Table S3) and analyzed the change in ubiquitylation at each site relative to the change in protein expression. This analysis demonstrated a range of stoichiometry changes. For some proteins, such as RSAD2 and IL1B, the increased quantity of ubiquitylated peptides was largely due to increased protein expression (Fig 3G). Conversely, proteins like IKBKE and GABARAP had large changes in ubiquitylation stoichiometry with minimal change in overall protein levels. Overall, changes in abundance of a ubiquitylated peptide and protein abundance were poorly correlated (R^2^=0.36) with changes in peptide abundance being driven by changes in stoichiometry of ubiquitylation in roughly 70% of cases.

We next examined ubiquitylation in CBL-deficient macrophages following *Mtb* infection. As expected, the loss of CBL did not change the abundance of most ubiquitylated peptides, consistent with a complex ubiquitin signaling landscape involving numerous E3 ligases. However, there was also a clear subset of peptides that underwent CBL-dependent ubiquitylation, including well-established CBL substrates such as the receptor tyrosine kinases MET and CSFR1. As anticipated, most changes seen in *Cbl*^−/−^ cells were peptides with decreased ubiquitylation; there were 430 ubiquitylation sites that decreased (Log2FC >1.0, FDR<0.05) and 90 sites increased in abundance (Fig 3H,I). Changes in the stoichiometry of ubiquitylation drove the changes in abundance of most ubiquitylated peptides, with very little correlation between changes in abundance for ubiquitylated peptides and protein expression (R^2^=0.09, Fig 3J).

Systematic analysis of the proteins with CBL-dependent ubiquitylation demonstrated the enrichment of several pathways and processes; some had clear immune-related functions such as regulation of phagocytosis (7.1-fold over-representation, p≤0.01), vacuolar acidification (13.7-fold, p=0.02), NFkB regulation (7.2-fold, p=0.02), and IFNGR signaling (4.2-fold, p≤0.01, Fig 3K). There were also factors known to be involved in the immune response to *Legionella pneumophila* (5.3-fold, p≤0.01), *Salmonella enterica* (3.2-fold, p ≤0.001, and *Mtb* itself (3.1-fold, p<0.01). Several other unanticipated pathways were also over-represented in the set of proteins with CBL-dependent ubiquitylation, including endoplasmic reticulum-associated protein degradation (ERAD) pathway (6.2-fold-fold over-representation, p≤0.001) and iron uptake (6.1-fold, p≤0.001). Taken together, these findings suggest that the set of proteins ubiquitylated by CBL extends far beyond its established role in growth-factor signaling at the plasma membrane and includes several other cellular processes throughout the cytoplasm and on membrane-bound organelles.

### Function of proteins with CBL-dependent ubiquitylation

We next sought to identify which proteins with CBL-dependent ubiquitylation played important roles during *Mtb* infection. We identified a subset of 43 proteins that had peptides with at least 80% reduction in ubiquitylation in *Cbl^−/−^* cells. Of these, 20 had annotated functions associated in some way with innate immunity (Fig 4A), and we analyzed this set of 20 proteins for CBL-dependent functions during *Mtb* infection. We hypothesized that if a protein is an important substrate of CBL, then loss of that protein should either phenocopy the CBL mutant (if positively regulated by CBL) or reverse the CBL phenotype (if inhibited by CBL). To analyze each of these factors we designed a lentiviral vector with two sgRNA cassettes, thereby allowing us to generate mutant CIM cell lines lacking either CBL or a putative CBL substrate, as well as double-mutant cells lacking both simultaneously (Fig 4B). Editing efficiency of >80% was confirmed by either TIDE analysis of genomic DNA (85), or by immunoblot of the targeted protein (Fig 4C).

**Fig 4.**
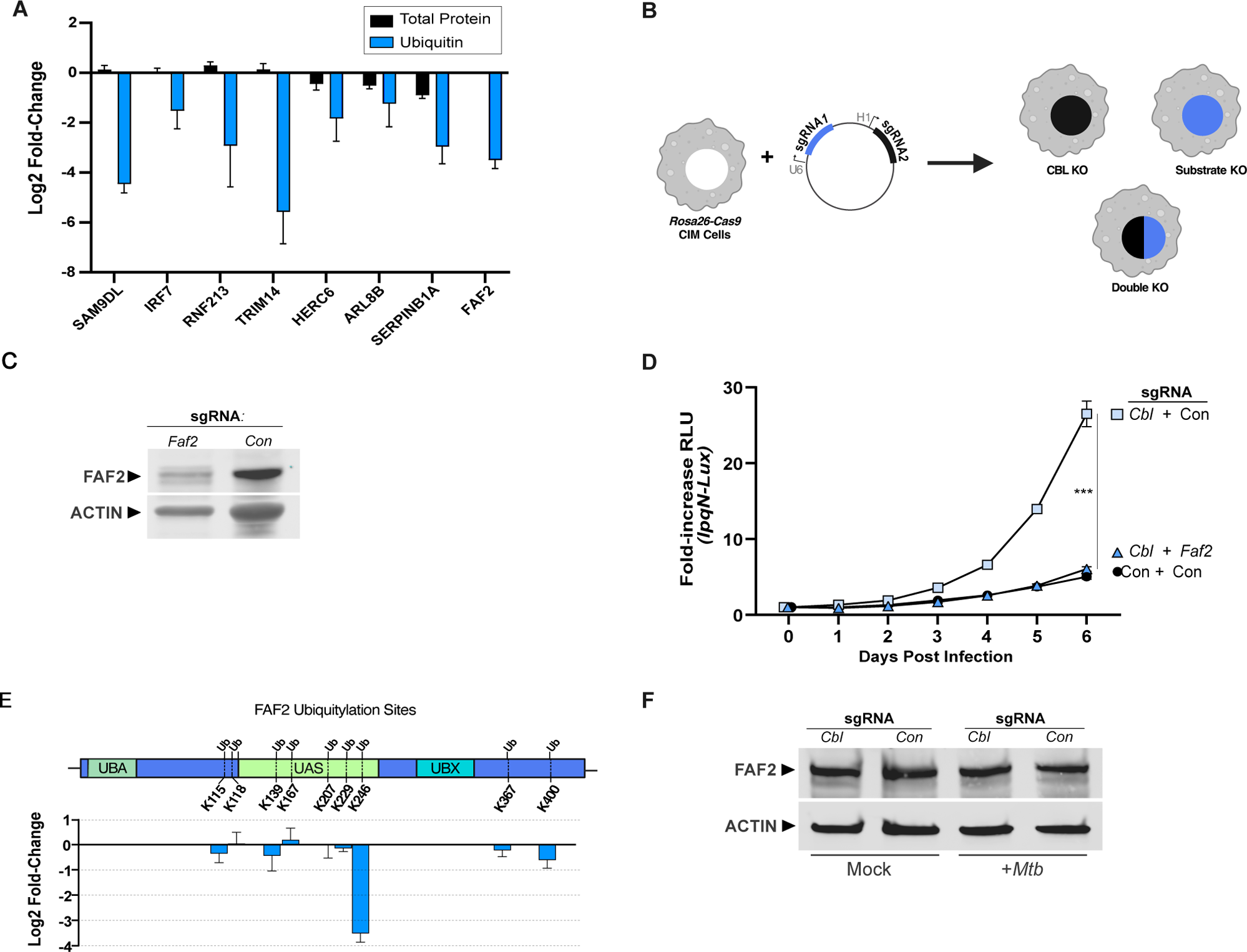
FAF2 promotes *Mtb* replication in the absence of CBL. (A) Examples of changes in ubiquitylation and total protein for immune regulators in *Mtb*-infected *Cbl*^−/−^ CIM cells 6 h post infection. In the case of proteins with multiple ubiquitylated peptides, the peptide with greatest statistically-significant fold-change is shown. (B) Experimental design for creation of cells lacking both *Cbl* and putative substrates. (C) Immunoblot of FAF2 protein in CIM cells expressing either non-targeting or *Faf2*-specific sgRNA. (D) Luminescent growth assay of CBL-sensitive *lpqN Mtb*. (E) Location of ubiquitylated residues on FAF2 and Log2 fold-change of ubiquitylation at each residue site in *Mtb*-infected *Cbl*^−/−^ cells vs. control cells. (F) Immunoblot of FAF2 protein levels in *Mtb*-infected control and *Cbl*^−/−^ cells. Error bars denote SEM of technical replicates; statistical significance was evaluated by two-tailed t-test. Representative (median) data of 4 independent experiments is shown.

We then infected each of these mutant cell lines with the CBL-sensitive *lpqN Mtb* mutant and assessed bacterial growth. As expected, the growth of *Mtb* was more rapid in CBL-deficient cells (Fig 4D), and in most cases, loss of a putative substrate in these CBL-deficient cells did not alter the replication of *Mtb* (data not shown). However, disruption of the Fas-associated factor 2 (*Faf2*) locus in CBL-deficient cells significantly altered the antibacterial capacity of these cells toward *Mtb*. FAF2 is an ER-localized signaling adapter protein, and our di-Gly MS analysis found it to be heavily ubiquitylated in macrophages, with 9 different ubiquitylated Lys residues. In *Cbl^−/−^*cells, a single residue (K246) showed a specific loss of ubiquitylation, with other Lys residues relatively unaffected (Fig 4E). In contrast to the other genes tested, disruption of FAF2 in CBL-deficient cells restored the ability of these cells to restrict *Mtb* replication, with bacterial growth rates approaching that seen in control macrophages that received a non-targeting sgRNA (Fig 4D).

Given the established role of CBL in targeting growth factor receptors for degradation, we tested the idea that CBL similarly targets FAF2 for degradation through ubiquitylation. However, FAF2 protein levels did not change following Mtb infection and were unaltered in CBL-deficient macrophages (Fig 4F). This suggests that rather than regulating FAF2 levels, CBL-dependent ubiquitylation is instead regulating the function of FAF2, perhaps by altering its protein-protein interactions. Taken together, these genetic analyses show that multiple proteins undergo CBL-dependent ubiquitylation, and FAF2 seems to be an important regulatory hub downstream of CBL as macrophages respond to *Mtb* infection.

## Discussion

### Antiviral versus antibacterial polarization

Macrophages must be able to sense and effectively respond to diverse types of pathogens. Both clinical data and animal models indicate that while a particular type of immune response may be beneficial in some circumstances, it can be detrimental in others. Tissue macrophages act as first responders in many infections, acting both to eliminate pathogens and to orchestrate subsequent immune responses. However, how a macrophage determines the type of pathogen it is encountering and translates that into an appropriate response remains unclear.

*Mtb* exposes an infected macrophage to an array of PAMPS; some are distinctively bacterially-derived molecules such as peptidoglycan or lipoarabinomannan, which activate NOD2 and TLR2 respectively. Activation of these sensors results in the release of cytokines with potent antibacterial activities such as TNF and IL1B (5–8,10,11). However, *Mtb*, as well as a number of other intracellular bacterial pathogens, including *Listeria monocytogenes,* and *Chlamydia trachomatis,* also activate cytosolic nucleic-acid sensors such as cGAS and induce expression of type I interferon and other antiviral effectors (63–66). While type I interferon potently inhibits viruses and can inhibit the growth of some bacteria such as *C. trachomatis*, it impairs antibacterial responses to *Mtb* and *L. monocytogenes* in mice (63–66).

The observation that antiviral responses can actively antagonize the antibacterial capacity of a macrophage suggests that macrophages might become polarized into antibacterial versus antiviral states. It also suggests that pathogens infecting macrophages like *Mtb,* and *L. monocytogenes* might seek to subvert this process as a virulence strategy. In the case of *Mtb*, it seems to do so through several mechanisms. First, upon perforating its phagosome, *Mtb* exposes bacterial dsDNA to the host cell cytoplasm which triggers cGAS-dependent activation of IRF3 and induction of IFN-β, as well as hundreds of other downstream genes, many of which have potent antiviral activity (24–31). Second, in parallel, *Mtb* also releases bacterial RNA into the host cytoplasm, thereby activating a similar RIG-I-dependent activation of IRF3, resulting in IFN-β expression (25). *Mtb* also introduces the LpqN virulence factor into host cells which interferes with the ability of the CBL ubiquitin ligase to regulate this process, thereby amplifying antiviral responses (45).

Exactly how the expression of antiviral effectors such as IFN-β antagonizes the antibacterial state of a macrophage is unclear. Integrating the results of different studies that examined different viral response regulators suggests that the transcriptional program controlling the expression of antiviral proteins such as IFN-β is likely to be comprised of several distinct modules, as the perturbation of different regulators causes distinct *Mtb*-related phenotypes. For example, in isolated ex vivo macrophages, disruption of CBL, TRIM14, or IRF7 alters the expression of antiviral effectors and alters the ability of a macrophage to restrict *Mtb* replication (45,46,67). In contrast, perturbation of RIG-I, SP140 or IFNAR has no effect on *Mtb* replication in isolated macrophages but profoundly alters *Mtb* susceptibility in mice (25,39,40,67–70). This suggests that while all these pathways regulate IFN-β signaling, they might differ in their regulation of other target genes, and that these differences drive distinct physiologic states in *Mtb*-infected macrophages. It also suggests that the additional regulatory inputs impinging on an infected macrophage during a complex immune response in vivo create a distinct environment where type I interferon signaling becomes a dominant factor.

### FAF2 Function

Although the mechanisms by which CBL regulates growth factor signaling are known in detail, the mechanisms by which it regulates innate immunity are not. CBL was initially characterized as a proto-oncogene that ubiquitylates activated receptor tyrosine kinases, thus targeting them for degradation, and was later found to have non-enzymatic functions in activating phosphatidylinositol 3-kinase and SRC-family kinases (48,49,51,71). We found that the enzymatic activity of CBL was needed for its ability to regulate immune responses during *Mtb* infection and identified the signaling adapter protein FAF2 as a critical intermediary factor, as disruption of FAF2 rescued the impaired responses in CBL-deficient cells. This suggests a scenario where the pathway regulated by FAF2 might somehow be hijacked by *Mtb* to impair host defenses, and that CBL normally acts to constrain FAF2 activity in *Mtb*-infected macrophages.

FAF2 is known to regulate a diverse set of physiologic processes, and thus the manner by which FAF2 regulates the innate immune response to *Mtb* is uncertain. FAF2 was originally characterized as a member of a family of ubiquitin regulatory X (UBX) domain proteins regulating the endoplasmic-reticulum-associated degradation pathway (ERAD) (72). FAF2 was also subsequently shown to act as a positive regulator of STING1, MTOR, and NFκB (73–75). Which of these pathways is the critical one downstream of FAF2 during *Mtb* infection is an important remaining question. It is also unclear what function the CBL-dependent ubiquitylation of FAF2 at K246 plays. A straightforward explanation would have been that CBL inhibits FAF2 by targeting it for degradation, analogous to the regulation of growth-factor receptors (49,76,77). However, our data demonstrate that this is not the case. Instead, it seems that ubiquitylation alters FAF2 functionality, perhaps by modulating its interactions with STING1, MTOR, or NFκB or their regulators. Future genetic experiments analyzing FAF2 alleles with mutations in K246 should provide insight into the function of FAF2 ubiquitylation, as will biochemical experiments analyzing the activation state of these different pathways.

### Ubiquitin signaling during *Mtb* infection

These studies also create a detailed map of protein ubiquitylation in macrophages as they respond to a pathogen, generating intriguing new hypotheses about how ubiquitin signaling might regulate innate immunity. Although numerous proteomics studies have analyzed ubiquitylation in mammalian cells, few of these focused on immune cells, and only a handful have examined signaling during innate immune responses to pathogens. Such studies in activated immune cells have provided important insights by identifying specific ubiquitylation events contributing to autophagy and DNA repair processes during infection that alter immune responses (78,79). Here, we identified ∼46,000 unique di-Gly-modified Lys residues, an in-depth dataset that significantly broadens the understanding of the ubiquitin-mediated signaling events taking place in macrophages as they respond to a pathogen. These analyses reveal a number of previously unknown ubiquitylation events on critical immune regulators, including the secreted cytokines IL1A and TNFSF13B, the endosomal peptidoglycan transporter SLC15A3, and the cytoplasmic peptidoglycan sensor NOD2 (Table S2). Thus, this dataset suggests multiple new mechanisms by which ubiquitin signaling might be regulating other critical immune responses.

### Limitations

While our RNA-Seq data show clear CBL-dependent transcriptional changes in CBL-depleted THP-1 macrophages, this is an imperfect model system. The expression of CBL is only reduced ∼85% by shRNA, and it is possible that residual CBL activity could be masking some changes. Although we also used CRISPR/Cas9 genome-editing to generate *CBL*^−/−^ THP-1 clones, we observed marked clone-to-clone variability in *Mtb*-related phenotypes making it an unsuitable alternative. In addition, the response of cell lines such as THP-1 cells to immune stimuli is often muted relative to primary macrophages (53). For example, we note that while changes in IFN-β expression were readily detectible in *Mtb*-infected THP-1 cells by RT-qPCR, the overall transcript levels were too low for reliable quantification by RNA-Seq. Separately, our MS studies had some limitations. They identified a large number of peptides with the di-Gly remnant, and while ∼95% of di-Gly Lys modifications result from ubiquitylation, NEDD8 and ISG15 both leave identical di-Gly remnants. Our current data are unable to distinguish the small subset of Lys residues modified by these other ubiquitin-related proteins. However, since CBL is believed to be a ubiquitin-specific ligase the set of CBL-dependent changes are likely due to ubiquitin. Finally, for those proteins with CBL-dependent ubiquitylation, we cannot distinguish at this time between direct ubiquitylation by CBL versus indirect effects mediated by another E3 ligase or deubiquitinase that CBL regulates, an analysis that will require complex in vitro ubiquitylation systems with MS analysis of reaction products, and important future direction.

## SUPPLEMENTAL FIGURE AND TABLE LEGENDS

**Fig S1.** (A) CFU determination in THP-1 cells expressing either CBL-specific or non-targeting control shRNA 5d after infection with *lpqN Mtb.* (B) Luminescent growth assay of *lpqN Mtb* in THP-1 cells depleted of CBL by a second independent shRNA.

**Fig S2.** (A) RNA-seq analysis of control THP-1 cells expressing a non-targeting shRNA, comparing uninfected cells and *lpqN Mtb-*infected cells. (B) RT-qPCR analysis of IFIT1 in IFN-β stimulated THP-1 cells.

**Table S1** RNA-Seq Dataset

**Table S2** Di-Gly MS Dataset

**Table S3** Total Protein MS dataset

## Materials & Methods

### Key Reagent Table

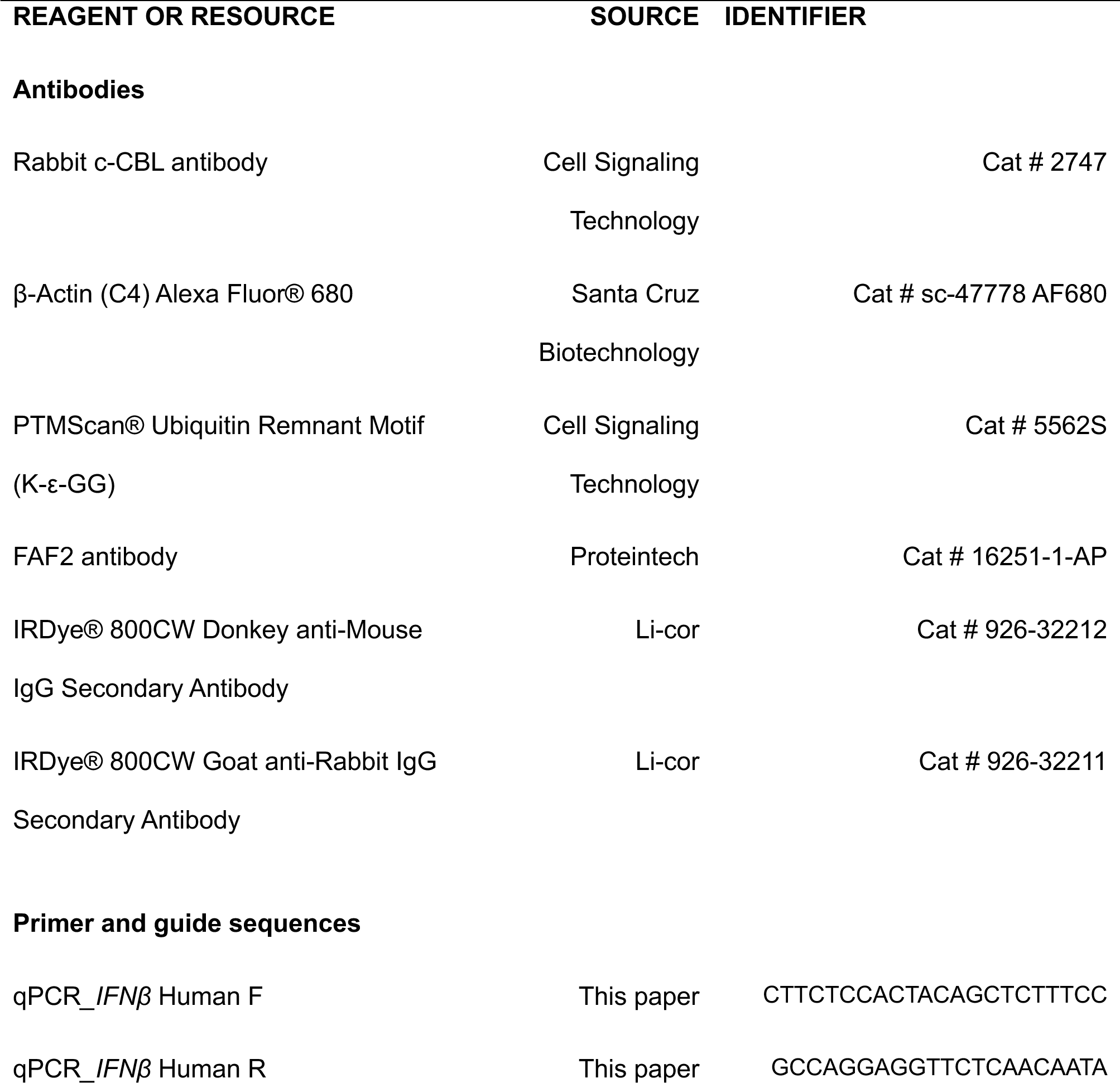
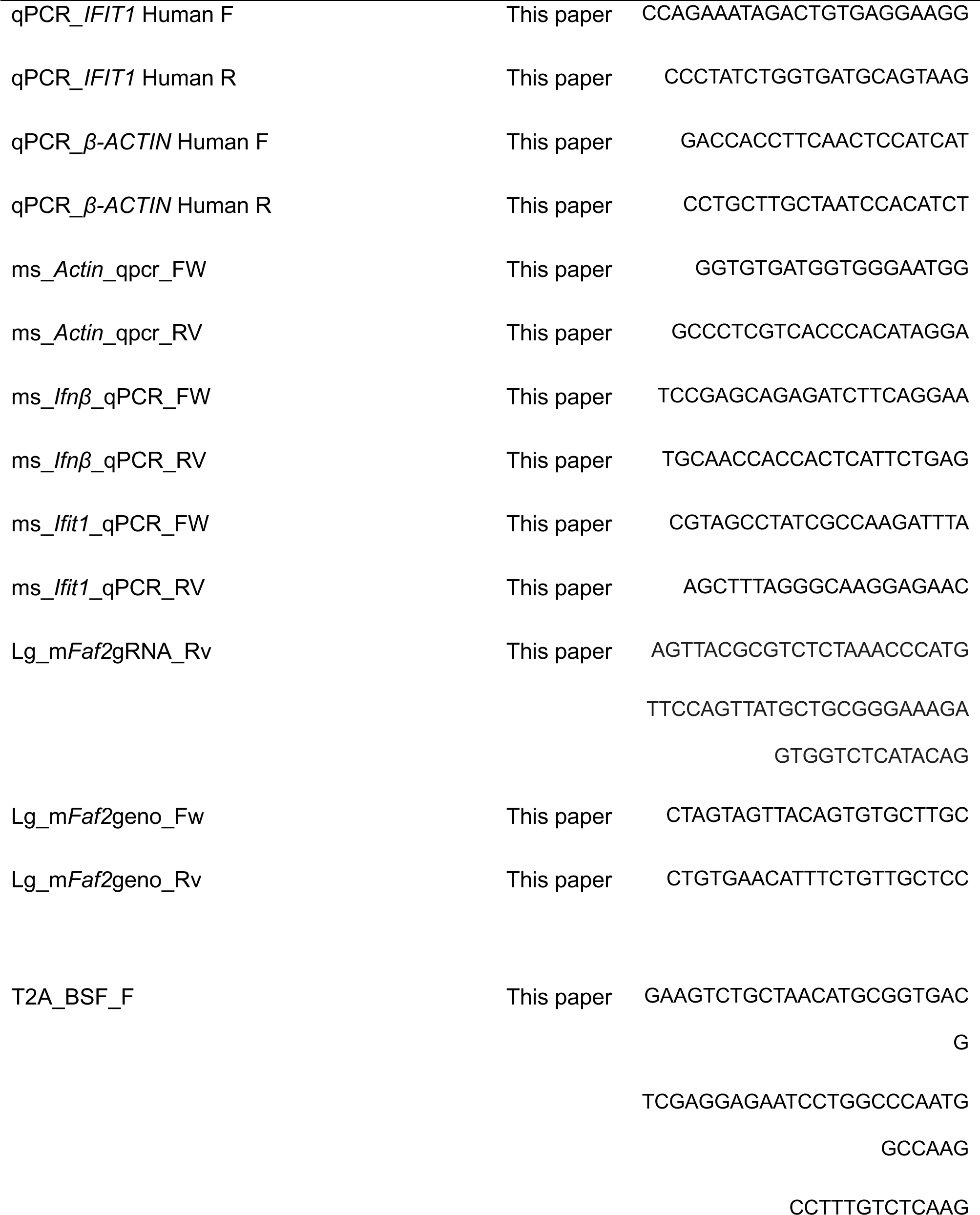
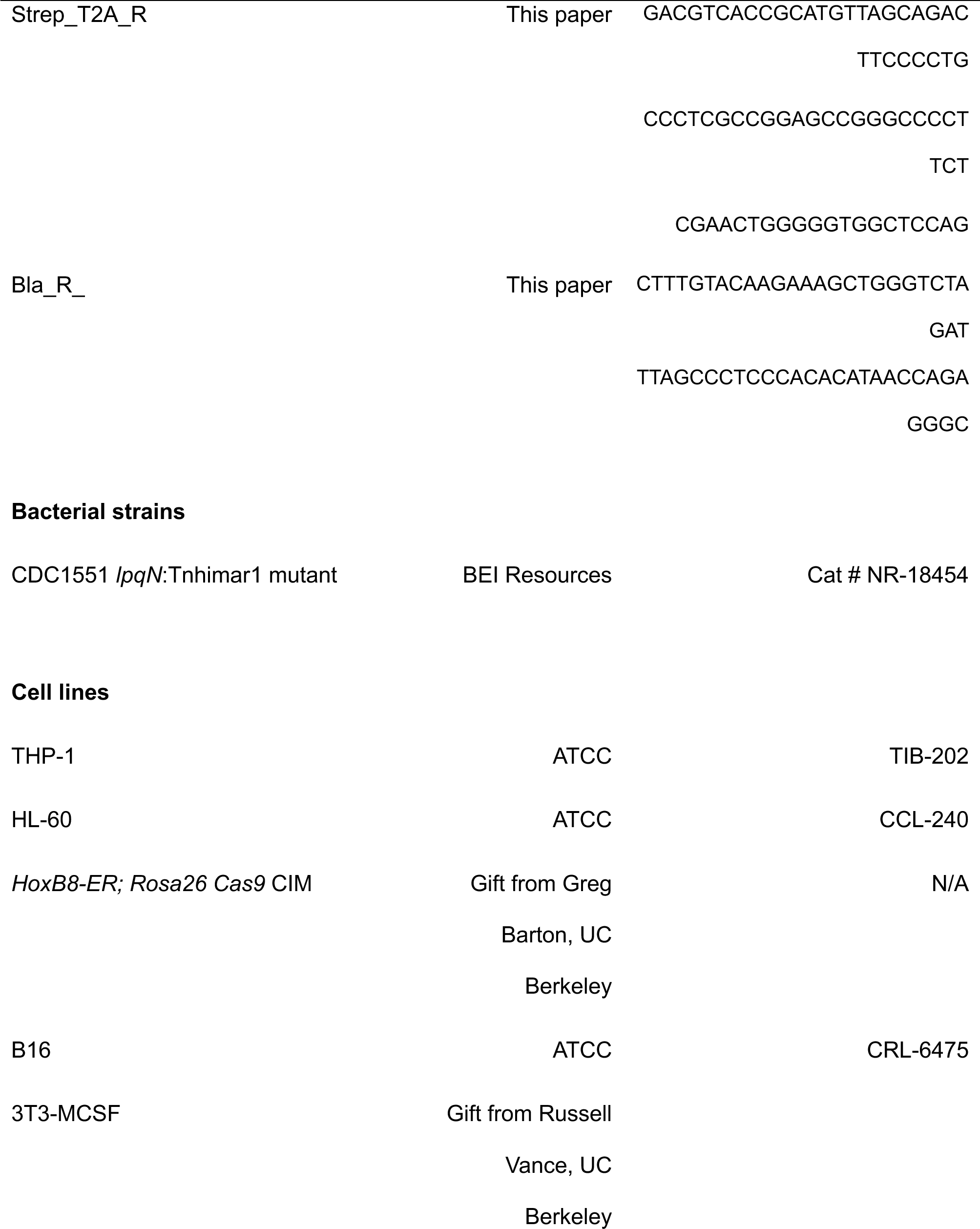
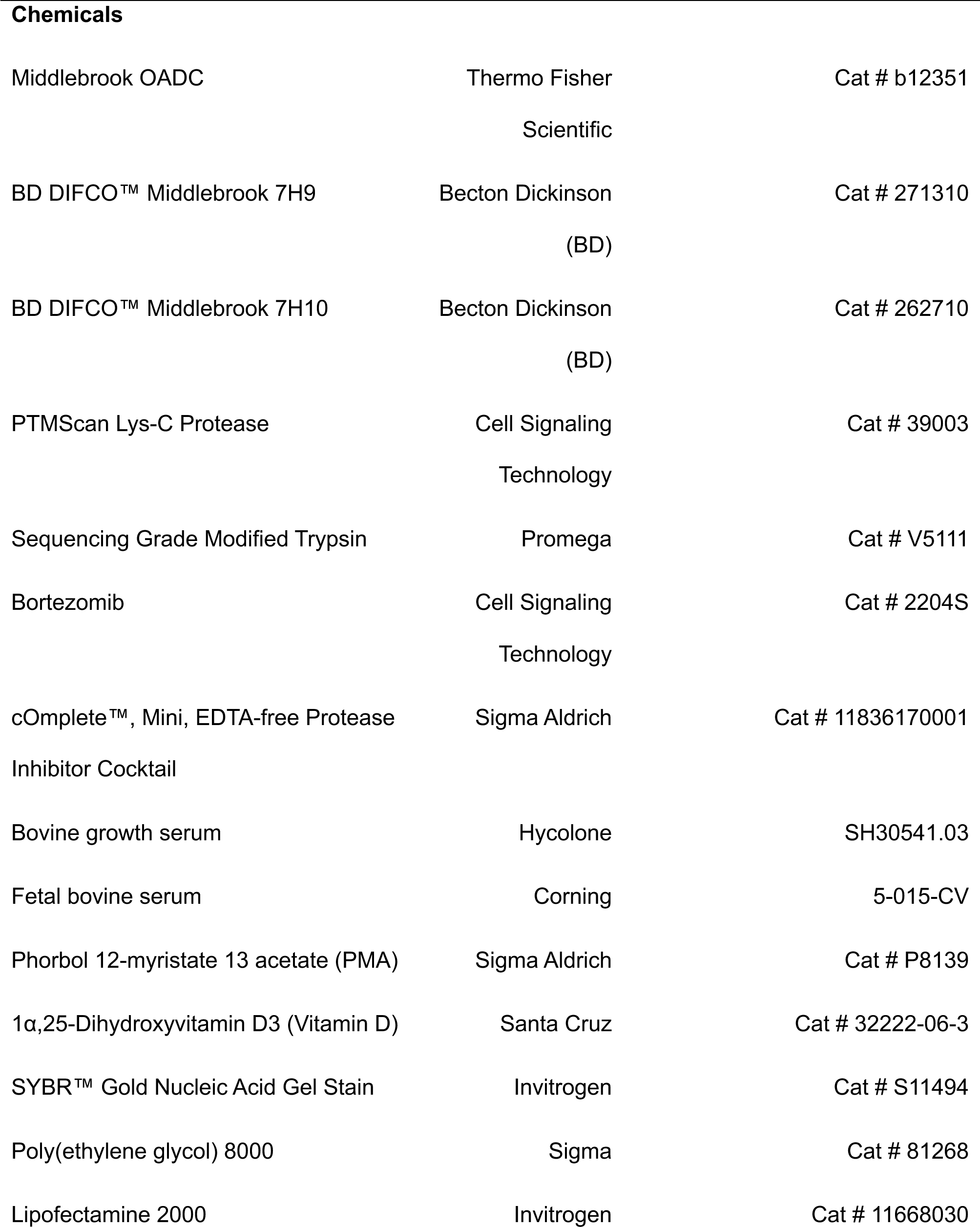
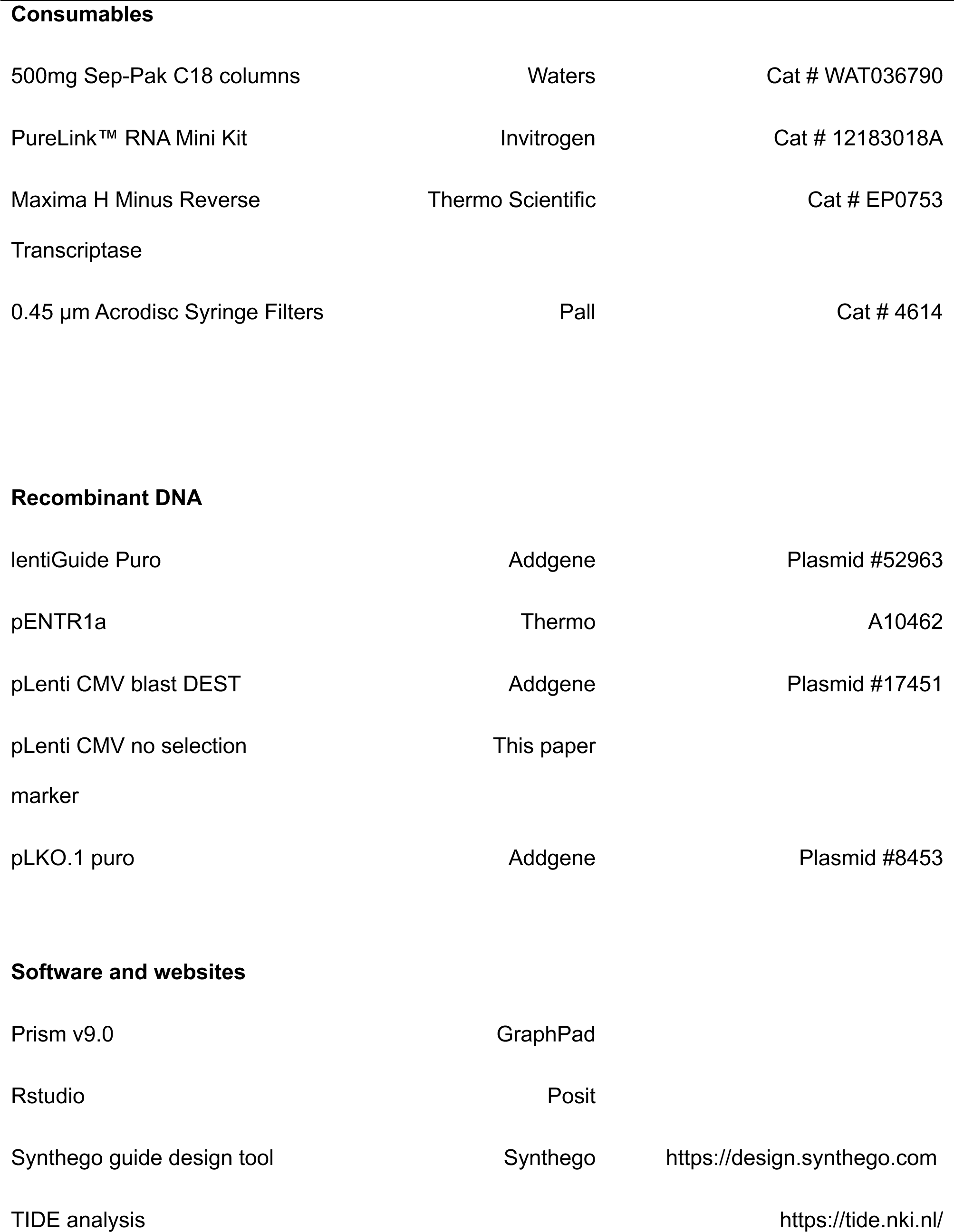
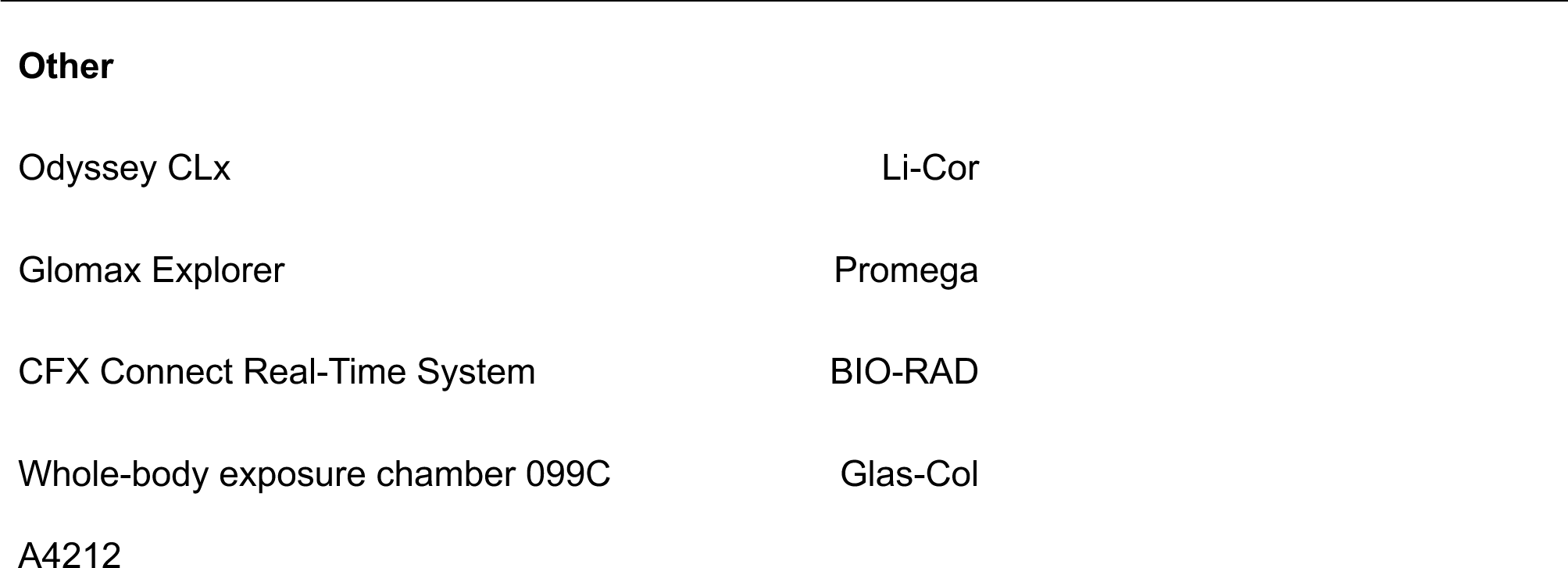

### Deposited data

At the time of publication raw MS data files will be deposited at MasIVE https://massive.ucsd.edu/ProteoSAFe. RNA-Seq data will be deposited at GEO https://www.ncbi.nlm.nih.gov /geo/info/seq.html. Processed MS and RNA-Seq data will be available as supplementary data files at the time of publication.

### Cell Lines

THP-1 human monocytes (ATCC TIB-202) and HL-60 promyeloblast cells (ATCC CCL-240) were purchased from ATCC. THP-1 cells were cultured in RPMI media containing 10% fetal bovine serum (FBS), 10mM HEPES, 1mM sodium pyruvate, 4500 mg/L glucose, and 0.05 mM 2-mercaptoethanol. HL-60 cells are cultured in RPMI containing 10% bovine growth serum (BGS), 15% fetal bovine serum (FBS), 1mM sodium pyruvate, 2mM L-glutamine, and 1X MEM non-essential amino acids solution. HOXB8-ER and Cas9-expressing conditionally immortalized macrophages (CIMs) were cultured in RPMI media containing 10% FBS, 10mM HEPES, 1mM sodium pyruvate, 2mM L-glutamine, 1.5ul 2-mercaptoethanol, 2% GM-CSF as conditioned media from B16 cells murine melanoma cells, and 2uM β-estradiol as previously described (53), prior to differentiation. THP-1 and HL-60 cells were differentiated with DMEM supplemented with 10% FBS, 1mM pyruvate, 2mM L-glutamine, 10ng/ml phorbol 12-myristate 13 acetate (PMA), and 0.1ng/ml 1,25 dihydroxy-vitamin D. Cells were differentiated for 72 h before experiments. CIM cells were differentiated by washing to removal of β-estradiol and then culture with DMEM supplemented with 10% FBS, 2mM L-glutamine, 1mM sodium pyruvate, 10% MCSF (derived from 3T3 MCSF conditioned media) for 7 days prior to experiments.

All cell lines generated using lentiviral vectors followed the same steps of transfection of HEK 293T Lenti-X cells with the transfer vector expressing the gene of interest, alongside the packaging plasmids psPAX2 and the envelope plasmid VSV-G. Viral supernatant is harvested 48 hours after transfection and either filtered with a 0.45 µm syringe filter and for CIM cell transduction, concentrated using A PEG-8000 solution (80). For transduction, viral supernatant with Polybrene 8ug/ml is added for 12h to THP-1 and HL-60 cells. CIM cells are transduced through spinfection and spun for 2h at 1000 RPM at 32C and then incubated for 6h before washing and culture in fresh media. Selective antibiotics were as follows: THP-1 and HL-60 cells are selected with 0.5 μg/ml of Puromycin or 4 μg/ml of Blasticidin. CIM cells are selected with 8 μg/ml of Puromycin or 4 μg/ml of Blasticidin.

### Bacterial Strains

*Mtb* was cultured in 7H9 (BD) supplemented with 0.5% glycerol, 10% oleic acid-albumin-dextrose-catalase (OADC, Middlebrook), and 0.1% Tween-80 in bottles gently rotated at 40 RPM at 37° C or on 7H10 plates supplemented with 10% OADC, 0.5% glycerol, and 0.1% Tween 80 at 37° C.

### Macrophage Infection for RNA harvest

*Mtb* strains were prepared for inoculation by washing twice in PBS with 10% heat-inactivated horse serum, gently sonicated, and spun at 500 rpm for 5 m to generate a fine bacterial suspension. Bacteria were opsonized in 10% heat-inactivated horse serum and macrophages were infected at an MOI = 10 for 6hr by spinfection at 1200RPM for 10 minutes followed by washing in PBS. After incubation, wells are washed once with 1X PBS and harvested in Trizol for RNA extraction.

### Macrophage Infection for Lux assay

For luminescent growth assays, cells are plated in triplicate in 96 well plates at a density of 8 x 10^4^ cells/well for THP-1 cells and 1.5 x 10^5^ cells/ well for CIM cells. Macrophage cell density at the time of infection was carefully matched between control and experimental cell lines to within <15% variance by plating sets of control wells across a gradient of densities and using SYBR gold to quantify nucleic acid content of plated experimental and control cells. Macrophages are infected at a MO1 = 0.8, and after spinfection plates are incubated for 30 minutes at 37° C. After the incubation, the monolayers were washed with PBS containing 1% heat-inactivated horse serum twice, and bacterial luminescence was measured over time in a GloMax Explorer warmed to 37 degrees C. Macrophage growth media was changed every other day. Colony-forming units (CFUs) were quantified by lysing triplicate wells of macrophages in sterile water and plated in serially dilutions on 7H10 agar with supplemented with OADC and 0.1% Tween-80. Plates were incubated at 37° C for 3 weeks prior to CFU enumeration.

### Western Blots

Protein in cell lysates was quantified by BCA. 30-50ug of protein lysate was separated by SDS-PAGE and transferred onto nitrocellulose membranes and blocked with LiCor blocking buffer at 0.5x for 1h. Primary antibodies were used at varying dilutions and incubated 1.5h at room-temperature or, in the case of anti-CBL antibody, 4° C overnight. Secondary antibodies were used at 1:10,000 dilution for 1h. After probing for indicated antibodies, the membranes were imaged on an Odyssey scanner (Licor).

### RNA Purification

Nucleic acid-stimulated macrophages were lysed in PureLink lysis buffer per manufacturer instructions. *Mtb-* infected macrophages were lysed in Trizol. In both cases, lysates were purified with silica spin columns (Purelink) per manufacturer instructions.

### Macrophage nucleic acid stimulation and RT-qPCR

Differentiated THP-1 and HL-60 cells are plated at 8 x 10^5^/ well in a 12-well plate. Cells are stimulated with 10ng of RNA or 20ng of DNA delivered by Lipofectamine 2000 as per manufacturer instructions and incubated for 6h. After RNA purification, cDNA was generated using 500ng total RNA with Maxima H minus reverse transcriptase and diluted 1:10 before qPCR analysis with the indicated primers and SYBR Green I detection of products.

### RNA-seq

Differentiated THP-1 cells were infected at MOI=10 for 6 h with *lpqN Mtb* in three independent experiments on different days. RNA was purified and barcoded 3’Tag-Seq libraries prepared using the QuantSeq FWD kit (Lexogen, Vienna, Austria) for multiplexed sequencing according to the recommendations of the manufacturer by the UC Davis DNA Sequencing core. The fragment size distribution of the libraries was verified via micro-capillary gel electrophoresis on a Bioanalyzer 2100 (Agilent, Santa Clara, CA). The libraries were quantified by fluorometry on a Qubit instrument (Life Technologies, Carlsbad, CA), and pooled in equimolar ratios. Twelve libraries were sequenced on one lane of an Aviti sequencer (Element Biosciences, San Diego, CA) with single-end 100 bp reads. The sequencing generated more than 4 million reads per library. Analysis of human data: HTStream was used to remove PhiX, adapter sequences, polyA tails, low quality sequences from end of reads, N bases, and reads less than 50 basepairs. After preprocessing, STAR was used to align reads to GRCh38.p13 human genome. UMI-tools was used to remove PCR duplicates post-alignment. Next, feature Counts was used to count reads whose alignment overlapped with genes using the gencode genome corresponding to the human genome version used. Annotation used was version 41. R was used for limma-voom pipeline on R with multiple testing correction using Benjamini-Hochberg procedure.

### Plasmid Construction for short shRNA and sgRNA delivery

shRNA expression plasmids were constructed by digesting pLKO.1 cloning vector with AgeI and EcoRI and annealed oligonucleotides containing the 21 shRNA encoding sequence were inserted, For sgRNA, 20 nucleotide targeting sequences were designed to target regions in the first exon of each locus using the Synthgo design tool. The targeting sequence is then fused by overlap-extension PCR to a second-generation tracRNA (81) and inserted into the HpaI and XhoI sites of pSicoR. The dual-guide sgRNA vector was constructed using a synthesized gene block containing the two guides of interest and an H1 promoter from Integrated DNA technologies (IDT) that were digested with BsmBI v2 on both ends and ligated to BsmBI v2 digested Lentiguide Puro vector (Addgene) with a second-generation tracRNA (81). Evaluation of editing efficiency was performed using TIDE.

### CRISPR/Cas9 genome editing

After lentiviral delivery of sgRNA cells were selected for 1 week with puromycin. The genome editing efficiency of the polyclonal population evaluated by PCR amplification of the targeted exon followed by Sanger sequencing and TIDE analysis. The polyclonal population was used for experiments.

### Site-directed mutagenesis

The wildtype and ligase mutant copies of CBL for both THP-1 and CIM cells contain synonymous mutations at the targeting sites of either the RNAi or CAS9 system to prevent degradation. Site-directed mutagenesis was carried out to generate catalytically inactive CBL mutants using pENTR1a plasmids (Addgene). Mutagenesis was validated through Sanger sequencing. Following validation, gateway cloning was used to move the mutated CBL open reading frame from pENTR1a into lentiviral pDEST vectors (Addgene). For THP-1 cells, constructs were cloned into a pLenti CMV Blast vector (Addgene). For CIM cells, constructs were fused by overlap extension PCR directly to a T2A peptide and blasticidin resistance gene and were cloned into pLenti CMV modified to remove other antibiotic resistance genes (82).

### di-Gly Enrichment

3 x 10^7^ cells were plated 24h before infection. Cells were infected with *lpqN Mtb* at an MOI = 10 for 6 hours. 2h prior to harvest, the 1μM of Bortezomib is added. For harvest, cells were washed 1X with PBS then scraped in ice-cold methanol+ 0.1 glycine pH 2.5 to precipitate protein, and inactivate proteases and *Mtb.* Cells were scrapped and removed from the BSL3. Chloroform was added to 20% volume to assist precipitation, incubated on ice for 10 min. and samples centrifuged to pellet protein. The pellets were washed once with ice-cold methanol, recentrifuged, and dried. Protein was resolubilized in in 9M Urea 25 mM HEPES pH 8.5,1 mM chloroacetamide (CAM), 0.5x Complete-mini protease inhibitor, and 0.5 mM EDTA with sonication. Proteins were then reduced for 45 min with 5 mM dithiothreitol (DTT) at room temperature and alkylated with 20mM CAM for 20 minutes at room temperature and in the dark. BCA assay was performed, samples diluted 1:4 in HEPES pH 8.0 and 2.5 mg of protein digested with Lys-C at enzyme to substrate ratio 1:100 at 37°C for 2h. Subsequently, samples were digested overnight at 37°C with trypsin enzyme to substrate ratio 1ug:100ug. Digestion was stopped with 0.5% TFA and then incubated on ice for 15 min to allow urea to precipitate and samples centrifuged to pellet debris. Peptides were desalted with 500 mg Sep-Pal C18 columns as per manufacturer instructions and lyophilized. 2.5mg of lyophilized peptides were resuspended with 0.5x immunoaffinity purification (IAP) buffer (50mM MOPS, pH7.2; 10mM sodium phosphate; 50mM NaCl) and sonicated. 10 ul cross-linked anti-K-ε-GG antibody beads were washed 3x in PBS. Samples were then mixed with 10 ul cross-linked anti-K-ε-GG antibody beads and incubated at 4°C for 2h. The peptide-bead mixture was then washed four times with PBS and one time with 0.1x IAP. Peptides were eluted with 100ul 0.15% TFA and analyzed by LC-MS/MS with on-tip clean up prior to MS run.

### Mass-Spectrometry

Chromatography was performed on an Evosep 1 using either the 60spd (di-Gly) or 30spd (total protein) method. Each sample was loaded onto a disposable Evotip C18 trap column (Evosep Biosytems, Denmark) as per the manufacturer’s instructions. Briefly, Evotips were wetted with 2-propanol, equilibrated with 0.1% formic acid, and then samples were loaded using centrifugal force at 1200g. Evotips were subsequently washed with 0.1% formic acid, and then 200 μL of 0.1% formic acid was added to each tip to prevent drying. The tipped samples were subjected to nanoLC on a Evosep One instrument (Evosep Biosystems). Tips were eluted directly onto a PepSep analytical column, dimensions: 150umx10cm C18 column (PepSep, Denmark) with 1.5 μm particle size (100 Å pores) (Bruker Daltronics), and a ZDV spray emitter (Bruker Daltronics). Mobile phases A and B were 100% water with 0.1% formic acid (v/v) and 100% Acetonitrile 0.1% formic acid (v/v), respectively. The standard pre-set method of 30 samples-per-day was used, which is a 44 min minute run or 60 spd which is a 21 min run.

Mass Spectrometry – Performed on a hybrid trapped ion mobility spectrometry-quadrupole time of flight mass spectrometer (timsTOF Pro 2 and timsTOF HT, (Bruker Daltonics, Bremen, Germany) with a modified nano-electrospray ion source (CaptiveSpray, Bruker Daltonics). In the experiments described here, the mass spectrometer was operated in diaPASEF mode. Desolvated ions entered the vacuum region through the glass capillary and deflected into the TIMS tunnel which is electrically separated into two parts (dual TIMS). Here, the first region is operated as an ion accumulation trap that primarily stores all ions entering the mass spectrometer, while the second part performs trapped ion mobility analysis.

DIA PASEF: The dual TIMS analyzer was operated at a fixed duty cycle close to 100% using equal accumulation and ramp times of 75 ms each. For the 30spd Evosep run, Data-independent analysis (DIA) scheme consisted of one MS scan followed by MSMS scans taken with 6×3= 18 precursor windows at width of 50Th per 6×3= 560 ms cycle, over the mass range 8×3= 300-1200 Dalton. The TIMS scans layer the doubly and triply charged peptides over an ion mobility −1/k0-range of 0.707-1.29 V*sec/cm2. The collision energy was ramped linearly as a function of the mobility from 59 eV at 1/K0=1.2 to 20 eV at 1/K0=0.7. For the 60spd Evosep run, Data-independent analysis (DIA) scheme consisted of one MS scan followed by MS/MS scans taken with 5×7= 35 precursor windows at width of 30^Th^ = 763 ms cycle time over the mass range 286-1307 Dalton. The TIMS scans layer the doubly and triply charged peptides over an ion mobility −1/k0-range of 0.7-1.3 V*sec/cm2. The collision energy was ramped linearly as a function of the mobility from 59 eV at 1/K0=1.2 to 20 eV at 1/K0=0.

### Data Analysis

LCMS files were processed with Spectronaut version 18.4(Biognosys, Zurich, Switzerland) using DirectDIA analysis mode. Mass tolerance/accuracy for precursor and fragment identification was set to default settings. The unreviewed FASTA for Mus Musculus, UP000000589, downloaded from Uniprot and a universal library of common laboratory contaminants (Frankenfield et al, 2022). For di-Gly analysis, a maximum of two missing cleavages were allowed, the required minimum peptide sequence length was 7 amino acids, and the peptide mass was limited to a maximum of 4600 Da. Carbamidomethylation of cysteine residues was set as a fixed modification, and methionine oxidation, acetylation of protein N termini and ubiquitination as variable modifications. A decoy false discovery rate (FDR) at less than 1% for peptide spectrum matches and protein group identifications was used for spectra filtering (Spectronaut default). Decoy database hits, proteins identified as potential contaminants, and proteins identified exclusively by one site modification were excluded from further analysis. For total protein: DIA data was analyzed using Spectronaut 18. using the direct DIA workflow with PTM localization selected. Trypsin/P Specific was set for the enzyme allowing two missed cleavages. Fixed Modifications were set for Carbamidomethyl, and variable modifications were set to Acetyl (Protein N-term), Oxidation, and ubiquitination. For DIA search identification, PSM and Protein Group FDR were set at 0.01%. Mass-spectrometry statistical analysis was performed in Spectronaut® (Biognosys) using t-test with multiple testing correction.

### Pathway enrichment analysis

Functional and pathway enrichment analysis for RNA-seq Ubiquitination Proteomics Gene ontology and pathway analysis were carried out on the set of genes with FDR <= 0.5 and LogFC <= 0.5 using the DAVID database (Database for Annotation, Visualization and Integrated Discovery; https://david.ncifcrf.gov/) (84). The significantly enriched biological pathways were identified with FDR <0.05. The pathways were visualized using R.

### Statistics

Statistical significance for RT-qPCR data was performed using GraphPad Prism version 10 (GraphPad Software, LLC) using two-tailed Student t-test when comparing. Statistical analysis for RNA-sequencing datasets was performed using the limma-voom pipeline on R with multiple testing corrections using Benjamini-Hochberg procedure. Ubiquitination statistical analysis was carried out within Spectronaut® (Biognosys) using the integrated statistical package with randomized imputation of values near the limit of detection if a peptide was undetectable in a particular sample, and performs t-tests with multiple testing correction and FDR calculation.

## Acknowledgements

We would like to thank Sebastian Winter and Jeroen Saeij for their input on this manuscript. We would like to thank the DNA Technologies and Bioinformatics cores at UC Davis that assisted with the sequencing and processing of the RNA-seq analysis. Schematic images were prepared with Biorender. BHP was funded by NIH 1R01AI144149.

