## Supplementary figures and images for "The balance between antiviral and antibacterial responses during *M. tuberculosis* infection is regulated by the ubiquitin ligase CBL"

### Fig S1

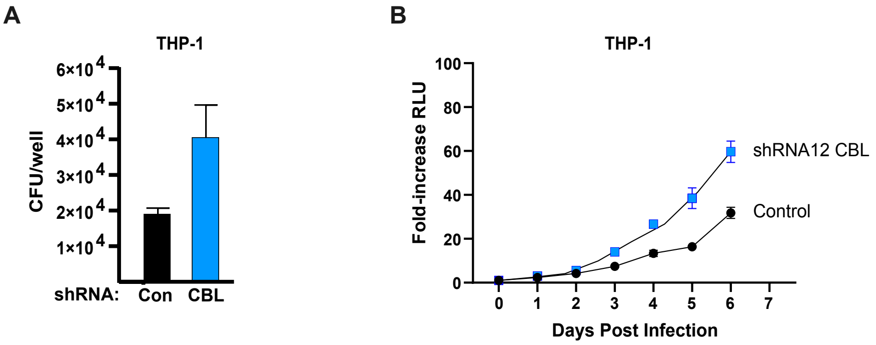

### Fig S2

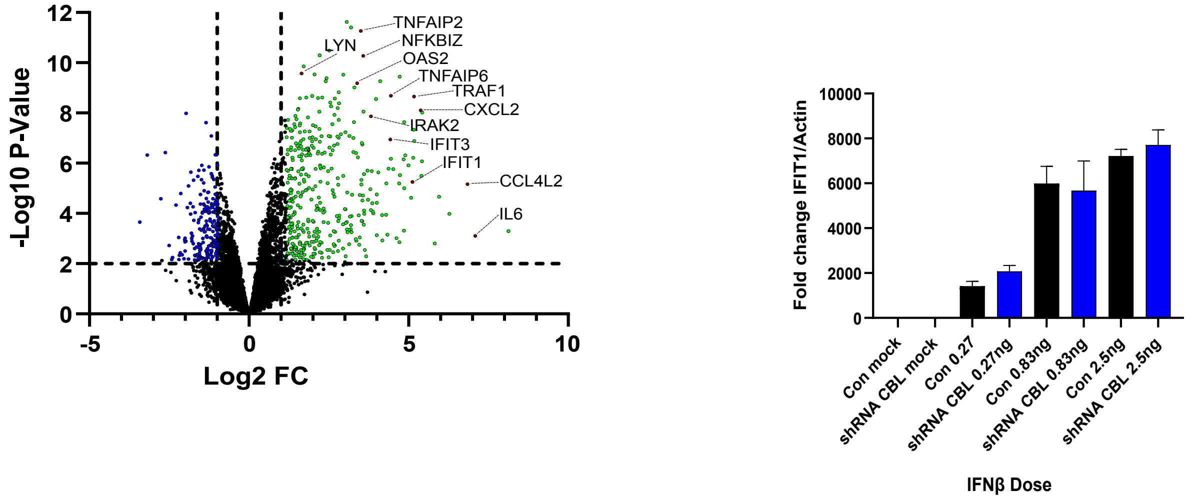
